# Trabeculations of the porcine and human cardiac ventricles are different in number but similar in total volume

**DOI:** 10.1101/2023.08.17.553743

**Authors:** Bjarke Jensen, Daniela Salvatori, Jacobine Schouten, Veronique M. F. Meijborg, Henrik Lauridsen, Peter Agger

**Affiliations:** Department of Medical Biology, Amsterdam Cardiovascular Sciences, University of Amsterdam, Amsterdam UMC, Amsterdam, The Netherlands; Department of Clinical Sciences, Anatomy and Physiology, Faculty of Veterinary Medicine, Utrecht University, Utrecht, The Netherlands; email DS,; JS; Department of Experimental Cardiology, University of Amsterdam, Amsterdam University Medical Centers, Amsterdam, the Netherlands; email; Department of Clinical Medicine, Aarhus University, Aarhus N, Denmark; email HL,; PA

**Keywords:** Heart, Development, Evolution

## Abstract

An intricate meshwork of trabeculations lines the luminal side of cardiac ventricles. Compaction, a developmental process, is thought to reduce trabeculations by adding them to the neighboring compact wall which is then enlarged. When pig, a plausible cardiac donor for xenotransplantation, is compared to human, the ventricular walls appear to have fewer trabeculations. We hypothesized the trabecular volume is proportionally smaller in pig than in human. Macroscopically, we observed in sixteen pig hearts that the ventricular walls harbor few but large trabeculations. Close inspection revealed a high number of tiny trabeculations, a few hundred, within the recesses of the large trabeculations. While tiny, these were still larger than embryonic trabeculations and even when considering their number, the total tally of trabeculations in pig was much fewer than in human. Volumetrics based on high-resolution MRI of additional six pig hearts compared to six human hearts, revealed the left ventricles were not significantly differently trabeculated (21.5 versus 22.8%, respectively), and the porcine right ventricles were only slightly less trabeculated (42.1 versus 49.3%, respectively). We then analyzed volumetrically ten pig embryonic hearts from gestational day 14 to 35. The trabecular and compact layer always grew, as did the intertrabecular recesses, in contrast to what compaction predicts. The proportions of the trabecular and compact layers changed substantially, nonetheless, due to differences in their growth rate rather than compaction. In conclusion, processes that affect the trabecular morphology do not necessarily affect the proportion of trabecular-to-compact myocardium and they are then distinct from compaction.

## Introduction

On the luminal side of mammal cardiac ventricles, there is a layer of myocardium that is organized in a trabecular meshwork [1, 2]. On the epicardial side, there is a wall of compact myocardium [3–6]. The trabeculations of the trabecular layer are too numerous to count macroscopically in human ventricles [7, 8]. In pig, the ventricular walls appear to consist of many fewer trabeculations [9]. This numerical difference presumably relates to the gestational process of compaction. In compaction, trabeculations coalesce into the compact wall whereby trabeculations become fewer in number while they add to and therefore thicken the compact wall [10].

Starting in the fifth week after conception in human, the embryonic heart forms numerous trabeculations [11]. Formation of new trabeculations has mostly ended a couple of weeks later [12]. In pig, a similar process is said to occur approximately in the third and fourth week of development [10]. These embryonic trabeculations will give rise to papillary muscles, trabeculations or trabeculae carneae, Purkinje cells, and, perhaps, compact wall in the fetal period [13–16]. If trabeculations are added to the compact wall, besides a reduction in their number, there should also be a reduction in the ventricular cavity in between trabeculations, the intertrabecular recesses [17]. Compaction, however, may be a subtle process only in mammals [18] not withstanding compaction seems to be a major driver of early ventricular septation in chicken and lizard [19, 20]. Rather than compaction, differences in growth rate of the trabecular and compact layer has a dramatic effect on the relative thicknesses of the two layers in human, mouse, and chicken [12, 21]. In addition, ventricular dilation may be an additional factor, at least in the postnatal heart, because it associates with an increase in the trabecular-to-compact ratio as seen in pregnant women and Duchenne’s muscle dystrophy [22, 23].

The notion of compaction as an important developmental process gained attention when it was hypothesized that its absence, so-called noncompaction, leads to a setting of excessive trabeculation and cardiomyopathy [24–29]. That noncompaction results in a greater-than-normal number of trabeculations in the fetus or adult, can be inferred from the following quotations: the “stratum spongiosum of each ventricle underwent differentiation but failed to resorb” [30]; the “gross anatomical appearance is characterized by numerous, excessively prominent trabeculations” [24]; and “altered structure of the myocardial wall as a result of intrauterine arrest of compaction of the myocardial fibers” [25]. Cohort studies and meta-analyses are now casting much doubt on the alleged association of excessive trabeculation and poor pump function [31–33]. In fact, a greater than normal extent of trabeculation has been associated with higher levels of physical activity [34, 35]. At tissue level, myocardium of the trabecular and compact layer are not different with regards to density of sarcomeres, vasculature, and mitochondria and the cardiomyocytes of the two layers do not differ in strength [21]. There is then a growing concern that the importance of compaction has been much exaggerated [18, 36].

Mice have been used much more extensively than pigs to study compaction [10, 37, 38]. The porcine heart, on the other hand, has the advantage of being similar in size to the human heart and the pig is now a realistic donor of hearts for transplantation to human patients [9, 39]. Because the anatomy and size of the trabecular layer may affect pump function, it is important to clarify how the porcine and human hearts differ in trabeculation. The trabecular layer can be quantified by the number or prominence of trabeculations, intricacy of the endocardial layer as assessed by a fractal dimension score, and the mass or width of the trabecular layer relative to the compact layer [24, 26, 40–46]. In this study, we investigated hearts in various states of contraction. Besides assessing the number of trabeculations macroscopically, we compared the absolute and relative volumes of trabecular and compact myocardium. Myocardial volumes, in principle, should be independent of the state of contraction, whereas the ratio of trabecular-to-compact wall thickness varies with the state of contraction [18]. The purpose of the study was to compare human and porcine hearts to test the hypothesis that the number of trabeculations in the adult ventricles predicts the proportion of trabecular and compact myocardium. Specifically, the prediction was that fewer trabeculations associate with a proportionally smaller trabecular layer.

## Materials and Methods

### Specimens

Sixteen pig hearts for the macroscopic investigations were collected at a local abattoir (in the Netherlands) where they had been excised from Dutch Landrace pigs of approximately 100 kg body mass that were slaughtered for human consumption. The protocols were consistent with the EC regulations 1069/2009 regarding slaughterhouse animal material for diagnosis and research as supervised by the Dutch Government (Dutch Ministry of Agriculture, Nature and Food Quality). The hearts were removed and immediately perfused with ice-cold cardioplegia (mM: 120 Na, 16 K, 1.2 Ca, 16 Mg, 144.4 Cl, 10 HCO_3_, 10 glucose, pH = 7.35–7.45) after which they were to be used in electrophysiological experiments. The hearts that eventually were not used were taken over by us and left in cardioplegia, unfixed, and refrigerated for approximately one week. Thereafter, rigor mortis had dissipated and the chambers walls were quite pliable while the macrostructure was intact. For the macroscopic inspections of the right ventricular luminal side (N=16), the free wall was resected by cuts immediately to the right of the interventricular sulcus. The left ventricular luminal side was exposed by a cut through the aortic root and valve, the fibrous continuity, septal leaflet of the mitral valve, and the mid-lateral wall to the apex (N=10).

The six porcine hearts for MRI originated from Danish landrace female swine each weighing 20 kg. The animals were pre-anesthetized with intramuscular administration of 0.5 mg/kg of midazolam and 2.5 mg/kg of ketaminol to allow establishment of an intravenous access followed by intravenous administration of 3 mg/kg of propofol to allow endotracheal intubation and coupling to a ventilator. Continuous anesthesia was maintained using 2–3% inhalational sevoflurane and analgesia was achieved with fentanyl (25 μg/kg/h). A median sternotomy was performed and after intravenous administration of 10.000 units of heparin, the animals were euthanized by means of exsanguination by excision of the heart.

Immediately after excision, the hearts were perfused with phosphate buffered formalin 10% (4% v/v formaldehyde) and stored in formalin for at least 48 hours. Twenty-four hours prior to MRI imaging, the hearts were perfused with phosphate buffered saline and were allowed to adapt to scanner room temperature. All procedures related to these six pigs were approved by the Danish Animal Inspectorate license no. 2013-15-2934-00869.

Eight human hearts were used and they came from the Becker archive at the Academic Medical Center, Amsterdam UMC. They were fixed in formalin in an unknown state of contraction. Two hearts were used for macroscopic inspection of the ventricular walls. Six hearts in which all chambers were intact were scanned with MRI.

Histological section series of whole pig embryos were used from archived material at the Faculty of Veterinary Medicine, Utrecht University. These sections were either 6, 8, or 10 µm thick and stained with hematoxylin-eosin. The section series came from embryos of gestational day 14 (series #13), 16 (series #14), 17 (series #15), 19 (series #35), 21 (series #18), 23 (series #20), 25 (series #22), 28 (series #25), 31 (series #27), and 35 (series #30). Between 13 (series #14) and 32 (series #18) equidistant sections were used to represent the whole heart (the median number of sections was 20).

### Histology of porcine ventricular wall

Transmural tissue blocks from various parts of the ventricular wall were imbedded in paraffin and cut in 10 µm sections. Sections were first stained yellow in Bouin’s fixative for one hour at 60 °C after which collagen was stained red in saturated picro-sirius red for one hour at room temperature followed by 2 min differentiation in 0.01 M HCl.

### MRI

Six human hearts were scanned in a Philips Achieva 1.5 T clinical system (Philips Medical Systems, Best, Netherlands), equipped with Nova Dual Gradients and Software Release 2.1.3. A 3D gradient echo imaging protocol was applied with a slice thickness of 0.5 mm and an in-plane-resolution of 0.5 × 0.5 mm^2^. Repetition time: 15 ms, echo time: 7 ms. Scan time was approximately 15 minutes for each heart. Six porcine and was scanned using an Agilent 9.4 T MR-system (Agilent, Santa Clara, California, USA), equipped with 400 mT gradients and vnmrJ 4.0 software. Anatomical data was gathered from the b0 images of a diffusion weighted sequence obtained using a standard multi-slice 2D spin echo sequence with diffusion gradients. The repetition time was 7000 ms and echo time was 30 ms. The scan time was approximately 16 h for each heart. A total of 125 slices with 800 μm slice thickness and no gap and was obtained with an in-plane-resolution of 0.4 × 0.4 mm^2^.

### Image analyses in Amira

Papillary muscles are sometimes excluded from clinical assessments of the trabecular layer volume. Nonetheless, because they derive from the embryonic trabeculations, like the remainder of the trabecular layer, the papillary muscles were included in the trabecular layer volume measurements [21, 47]. Image stacks were imported to Amira (v2020.2, Thermo-Fisher). On equidistant images in the transverse plane of the heart, we labeled the trabeculations and compact wall of the ventricular walls for both left and right ventricles in addition to the walls of the atria. The same approach was used for the embryonic porcine hearts. There is no overt boundary between the trabecular and compact layers, neither on thick sections nor on histology (while, a contrary example is seen in some species of fish [48]). Here, we strove to set the trabecular-compact boundary by connecting the deepest parts of the intertrabecular recesses. The detection of trabeculations, and the intertrabecular recesses, is dependent on spatial resolution [8], and we cannot establish whether the spatial resolution used here allows for the detection of the smallest and deepest parts of the recesses. Approximately 10 images were labeled for each chamber.

According to Cavalieri’s principle [49] or Simpson’s rule, these relate to the true volume with an error of approximately 10 %. Volume readouts of the labels were derived from the Materials Statistics module in Amira. They were then multiplied by the fixed distance between the labeled images.

### Statistics

Measured volumes between two chambers were compared using two-tailed paired t-tests without the assumption of equal variance. The relative trabecular volume of the left and right ventricles were compared with a linear regression to test whether greater trabeculation of one chamber associates with greater trabeculation of the other chamber. A p-value less than 0.05 was considered significant. All statistics were done in Excel (version 16.16.27, Microsoft, USA).

## Results

### Gross morphology of the right ventricular wall in pig and human

The right ventricle free walls of pig and human are the same in the sense that trabeculations cover almost the entire inner surface (Figure 1). Only the wall most proximal to the pulmonary valve is without trabeculations. This a-trabecular region was never large and varied between approximately 2-6 % of the total RV free wall. Compared with the human heart, the RV wall of the porcine heart has much fewer trabeculations (Figure 1). Most of the trabeculations, however, are large and if they are evaluated by size, many of them compares to the human papillary muscle (see labels 1-5 in Figure 1). Macroscopically, they may pass for ‘false pillars’ since most of them are fused with the compact wall and a probe cannot be passed behind them. In the human RV, by contrast, numerous trabeculations connect with neighboring trabeculations and these are then individual struts in a meshwork. In pig, free-standing trabeculations are mostly confined to the areas around the parietal papillary muscle group and the wall most proximal to the hinge line of the tricuspid valve.

**Figure 1.**
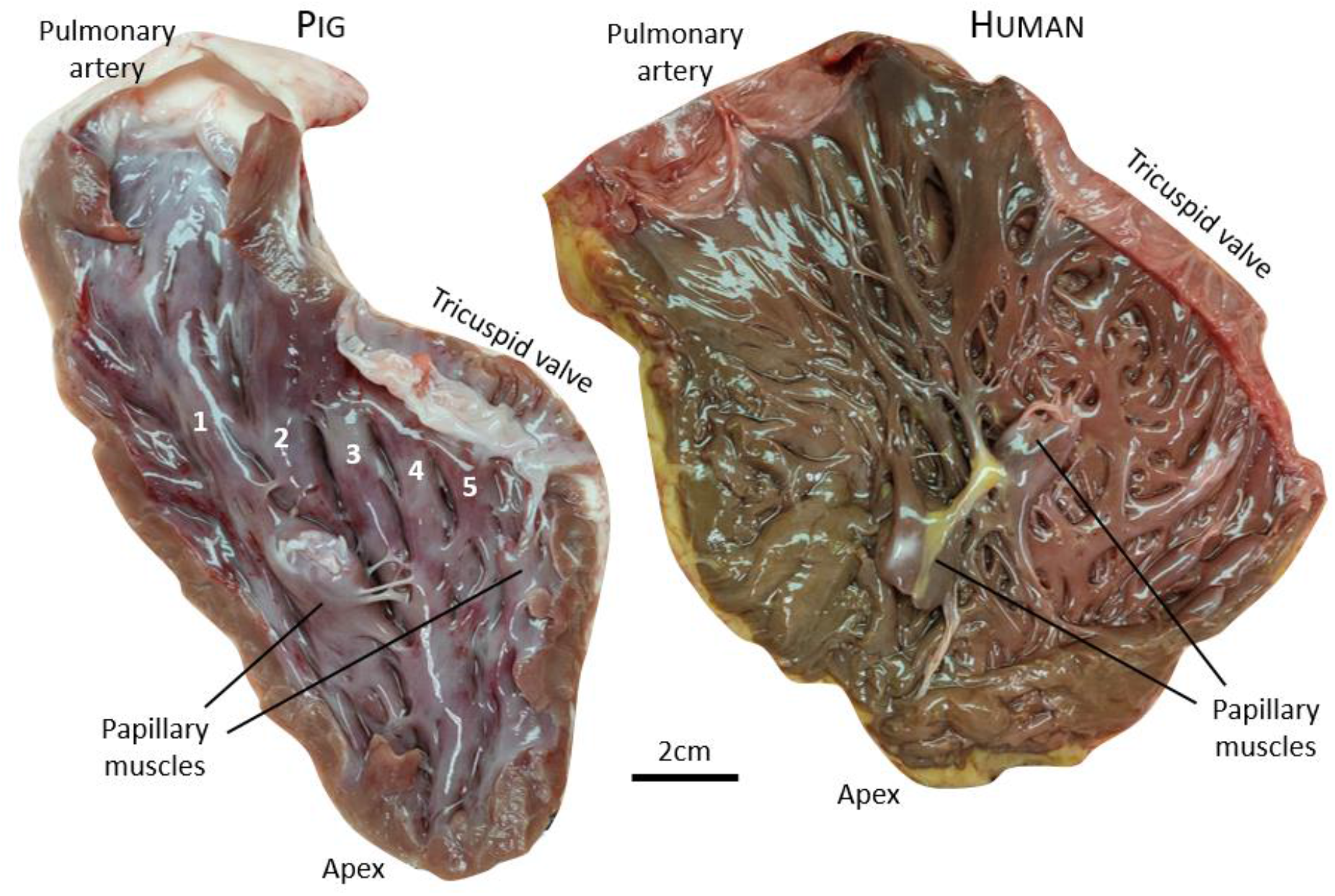
The right ventricular free wall of pig and human. The same principal components are present in both ventricles (valves and papillary muscles) and they have approximately the same position relation to each other in both species. The porcine wall, however, is dominated by a few very large trabeculations, five of which are indicated with numbers 1-5. In contrast, the trabecular layer of the human right ventricle appears much more as a meshwork of a high number of small trabeculations. In both specimens, the tendinous cords of the tricuspid valve have been cut and the leaflets are pulled toward the right atrium to fully expose the trabeculations.

**Table 1.** Characteristics of the six human hearts that were scanned with MRI.

| Case | Year of death | Age | Sex | Medical history | Cardiac pathologies (on MRI) | Coagulates |
| --- | --- | --- | --- | --- | --- | --- |
| S77_076 | 1977 | 10 | F | Bicuspid PAV | Possibly small membranous VSD | No |
| S86_042 | 1986 | 59 | M | Lung cancer | - | No |
| S96_166 | 1996 | 69 | F | ALS, RF | Potential PLSCV | No |
| S2000_43 | 2000 | 86 | F | ALS, SD | PFO | RA, RV |
| S2000_98 | 2000 | 70 | F | slow progressive ALS | LV scar/infarct lateral apical region<br>RV much fatty infiltration | LA, LV<br>RA, RV |
| T93_6067 | 1993 | ? | ? | ? | Aortic root dilation | RA, RV |

Notes. ALS, amyotrophic lateral sclerosis; F, female; LA, left atrium; LV, left ventricle; M, male; PAV, pulmonary arterial valve; PFO, persistent foramen ovale; PLSCV, persistent left superior caval vein; RA, right atrium; Respiratory failure; RV, right ventricle; SD, sudden death; VSD, ventricular septal defect; ?, unknown.

### Variation in macroscopic trabeculation of the right ventricle in pig

We noted that the trabecular layer of the porcine right ventricle was never fully identical between hearts. For example, the trabeculations immediately below the tricuspid valve in the free wall typically takes the form of approximately four to six approximately even-sized trabeculations with a long-axis from the hinge line of the tricuspid valve towards the ventricular apex (Figure 2A). These trabeculations may be fused just below the hinge line, and in one out of six hearts this fusion assumed the appearance of a transverse trabeculation (Figure 2B). The ventral-most two trabeculations or so will reach the base of the papillary muscle whereas the dorsal-most trabeculations will reach deeper towards the apex. In four out of six hearts the trabeculations reached the apex (Figures 2C), whereas in the remaining two hearts the trabeculations were quite coalesced, the intertrabecular recesses were shallow, and the overall appearance in the apical region was a sort of plateau (Figure 2A). The papillary muscle was positioned similarly in all six hearts, half-way down and immediately to the right of a line extending from the apex to ventral-most hinge of the tricuspid valve (Figure 2A). It was a solitary structure in five hearts, in the sixth heart there was a smaller second papillary muscle juxtaposed dorsally to the main one. There was much variation in how the moderator band connected to the papillary muscles; in two hearts there was a thick connection (Figure 2A), in two hearts only a thin strand, and in the remaining two hearts the moderator band connected into the trabeculations of the free wall ventrally to the papillary muscle (Figures 2D). From the apex to the base of the pulmonary valve, most trabeculations appeared oriented along this axis. Just below the pulmonary valve, the wall appeared a-trabecular, but this region (*pars glabra*) is always small (Figure 2A) if not very small (Figure 2E). Consequently, almost the entire cavity of the right ventricle can be considered lined with trabeculations.

**Figure 2.**
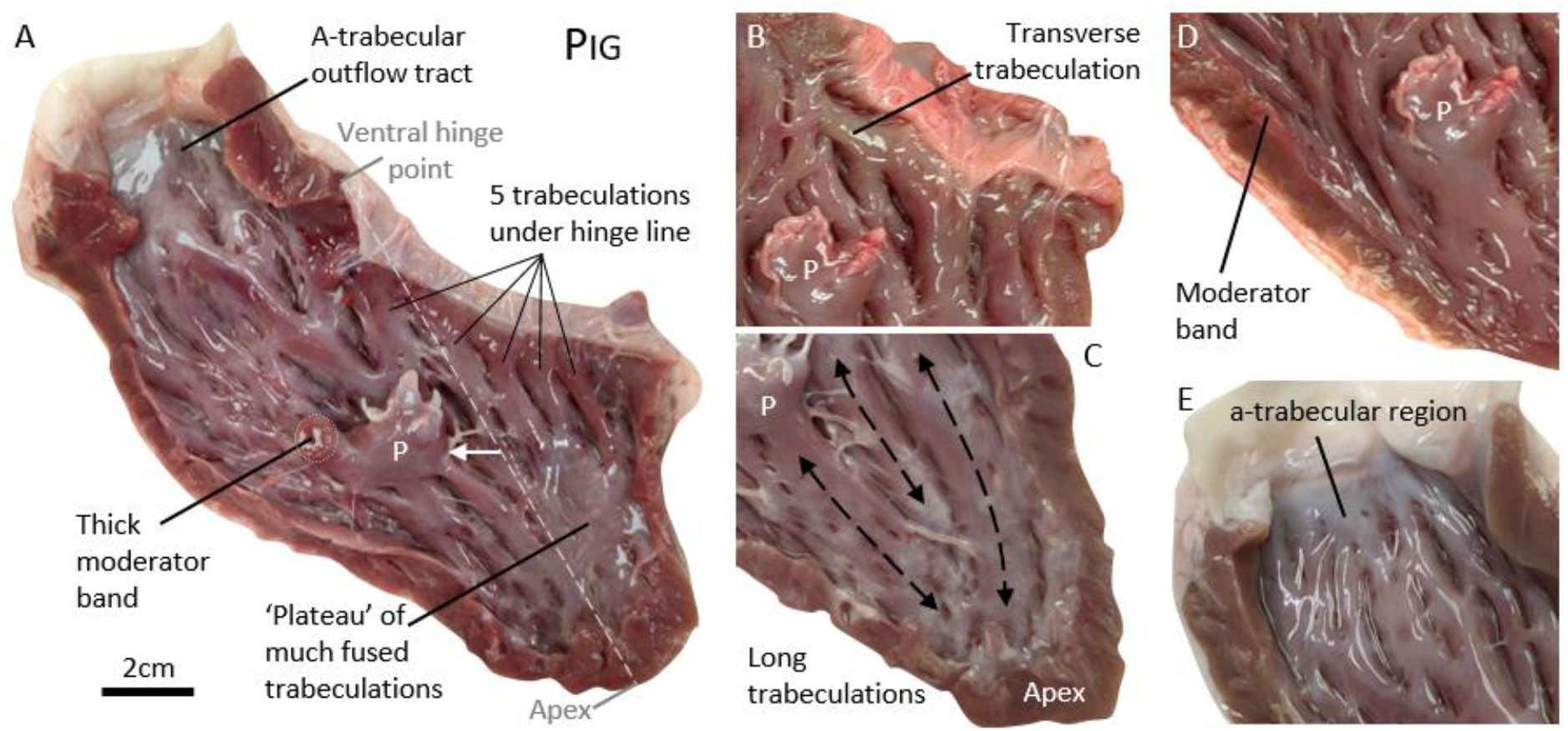
Variation in macroscopic trabeculation of the right ventricle. **A**. Overview of the porcine right ventricular free wall. **B**. A large transverse trabeculation was found under the hinge line of the tricuspid valve in one heart. **C**. A few large and long trabeculations reached the apex in four hearts, whereas these were fused into a plateau in two heart as exemplified in A. **D**. In two hearts the moderator band would connect to the free wall rather than to the papillary muscles as exemplified in A. **E**. Compared to the total wall, the a-trabecular region of the outflow tract could be a very small fraction.

### Small trabeculations in the porcine and human right ventricle

In the human right ventricle, there are numerous free-standing trabeculations and this is particularly apparent between the papillary muscles and the compact wall (Figure 3A). A similar setting is found in pig, only the trabeculations are fewer than in human (Figure 3B). In pig, these trabeculations are much smaller than the large trabeculations that dominate the macroscopic appearance of the free wall. Generally, small trabeculations are prevalent in the deep recesses between the large trabeculations (Figure 3C) and many of these likely remain obscure even under close inspection of deep recesses. These trabeculations were myocardial (Figure 3D-H) and despite their diminutive appearance, they were substantially larger than trabeculations of embryos. We did not investigate whether they contain Purkinje cells, but either way their origin is most likely embryonic trabeculations. The solitary small trabeculation in Figure 3F, for example, is longer (700 µm) and wider (300 µm) by approximately one order of magnitude than an embryonic trabeculation. Even the tiniest trabeculations we saw were larger than embryonic trabeculations (see the struts in the enclosed area of Figure 3G). A precise tally of all large and small trabeculations, then, was not possible. In addition, numerous dimples could be found, and it could not be determined whether these were unrelated to trabeculations or whether they were small and shallow intertrabecular recesses (Figure 3B). These difficulties of counting trabeculations were prevailing in the human right ventricle. Within the encircled area in Figure 3A, for example, some 25 semi-superficial trabeculations can be seen but there are undoubtedly more trabeculations deeper in the trabecular layer (in Figure 3B, a similar count yields some 15 trabeculations). Considering the entire free wall, a cautious estimate could be that in pig there are approximately 200 trabeculations whereas in human there could easily be more than 1000.

**Figure 3.**
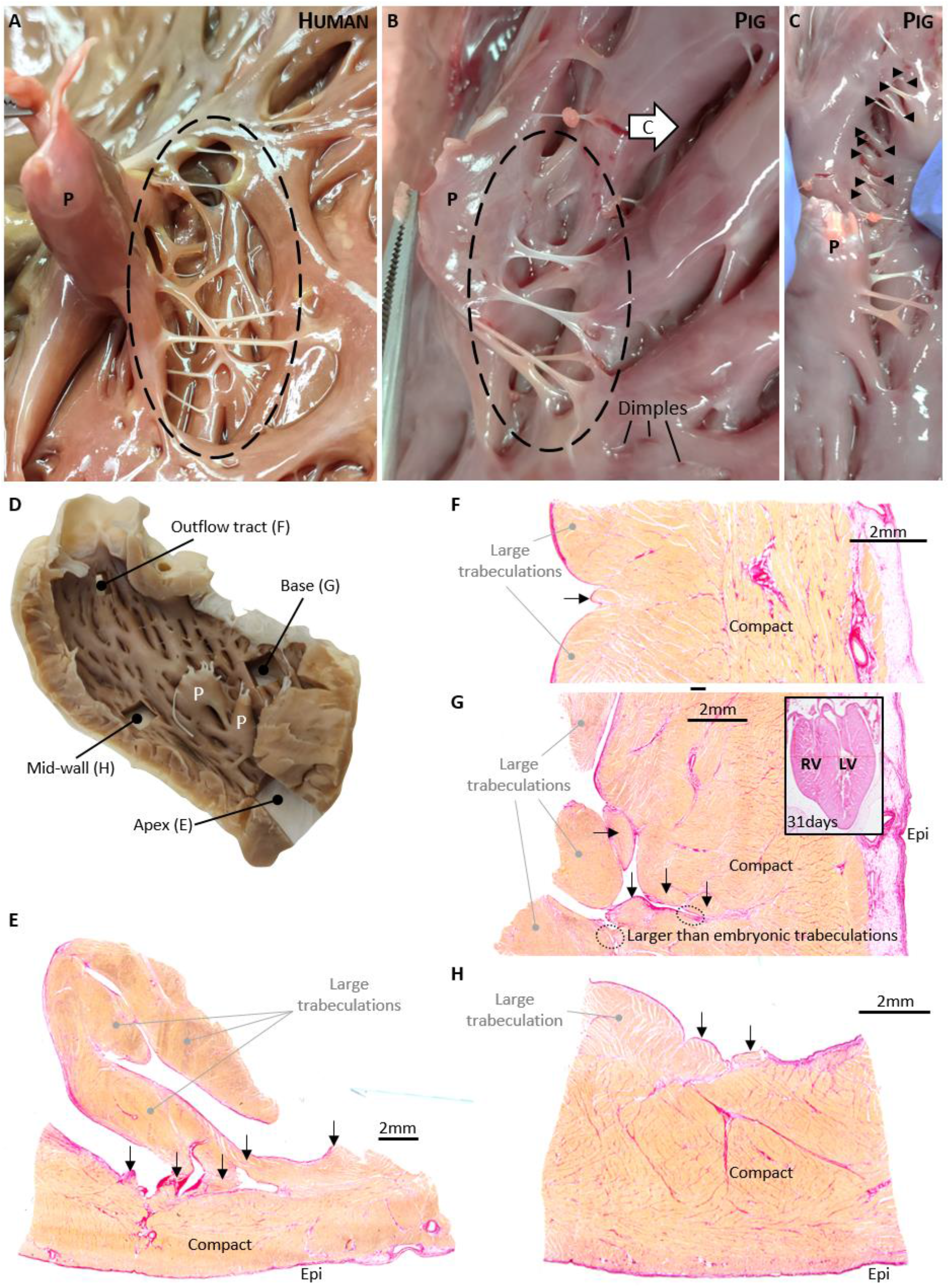
Small trabeculations of the right ventricle. **A**. Human right ventricle, showing approximately 25 trabeculations in the region marked by the dashed line which is between a papillary muscle (P) and the compact wall. **B**. Pig right ventricle, similar region as in A, showing approximately 15 small trabeculations in the region marked by the dashed line. The arrow marked with C, indicates the line of inspection of the intertrabecular recess in image C. **C**. Two large trabeculations are pulled apart which exposes approximately 12 small trabeculations. **D**. Overview of porcine right ventricle free wall and four sample sites for histology (stained with picro-sirius, red is collagen, orange is myocardium). **E**-**H**. Small myocardial trabeculations (arrows) revealed by histology of the apex (**E**), outflow tract (**F**), base (**G**), and mid-wall (**H**). In **G**, notice the tiny myocardial struts in the enclosed areas, which, despite their diminutive size are substantially larger than embryonic trabeculations (the insert is a 4-chamber view of the pig heart at gestational day 31 and both ventricles contain a high number of tiny trabeculations).

### Gross morphology of the left ventricular wall in pig and human

The left ventricle free wall of pig and human are similar in that trabeculations cover almost all of the inner surface of the chamber wall (Figure 4). Trabeculations are only absent in the septal part most proximal to the aortic valve, the outflow tract region (*pars glabra*). Compared with the human heart, the LV wall of pig heart has much fewer trabeculations (Figure 4). Most of the trabeculations are large and by size, many of them could almost pass for a human papillary muscle. Of these, there are typically four particularly large trabeculations with an apex-to-base orientation on the septal surface between the anterior and posterior papillary muscles (N=3, see labels 1-4 of Figure 4). The porcine papillary muscles are very big. Like in the RV, most of the porcine large trabeculations and the base of the papillary muscles are fused with the compact wall so that a probe cannot be passed behind them. In the human LV, by contrast, the papillary muscles have a trabecular base [50] and numerous trabeculations connect to neighboring trabeculations and these are then individual struts in a meshwork (Figure 4).

**Figure 4.**
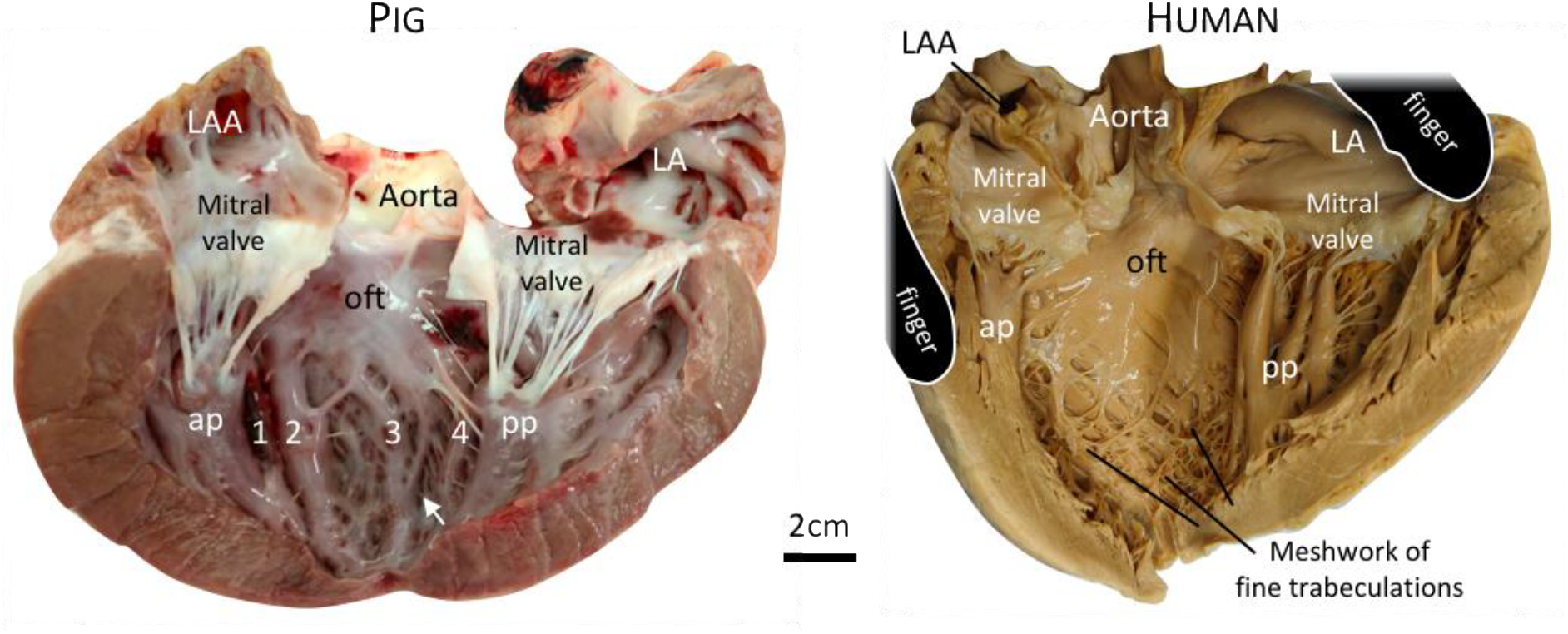
Left ventricle of pig and human. The same principal components are present in the ventricles of pig and human - mitral valve, anterior (ap) and posterior papillary muscles (pp), outflow tract (oft) - and they have approximately the same position in relation to each other. In addition, the porcine wall is dominated by a few very large trabeculations, four of which are indicated with numbers 1-4, whereas in the human left ventricle there is a meshwork of a much higher number of small trabeculations. Small trabeculations can also be found in the porcine left ventricle, one example is indicated by the white arrow, but these are much fewer than in the human heart. In both specimens, the fibrous continuity of the mitral valve to the aortic root has been cut to expose the outflow tract.

### Myocardial volumes of the porcine and human hearts

The volume of myocardium of the pig heart was estimated based on labelling of equidistant MRI images of the transverse plane (Figure 5A-B). On average, the total myocardial volume (atrial and ventricular myocardium combined) was 94.7 ml (±7.1, standard deviation). As expected, the left ventricle had the most myocardium, followed by the right ventricle, and the volume of the left and right atrial walls were smaller than that of the right ventricle (Figure 5B). The fraction of atrial to ventricular myocardium was 0.15 (±0.02, SD), and thus typical of vertebrates [51], while the fraction of right atrium to left atrium was 1.01 (±0.18, SD) and not different from 1 (one sample t-test for difference from 1, P=0.931). Both left and right ventricle had more compact than trabecular myocardium (Figure 5C) and the left ventricle was proportionally significantly less trabeculated than was the right ventricle (21.5% ± 1.7 versus 42.1% ± 1.4, paired t-test, P<0.001). Finally, using a Pearson correlation, we found that the proportion of trabeculation of the left ventricle was not correlated to that of the right ventricle (P=0.583).

**Figure 5.**
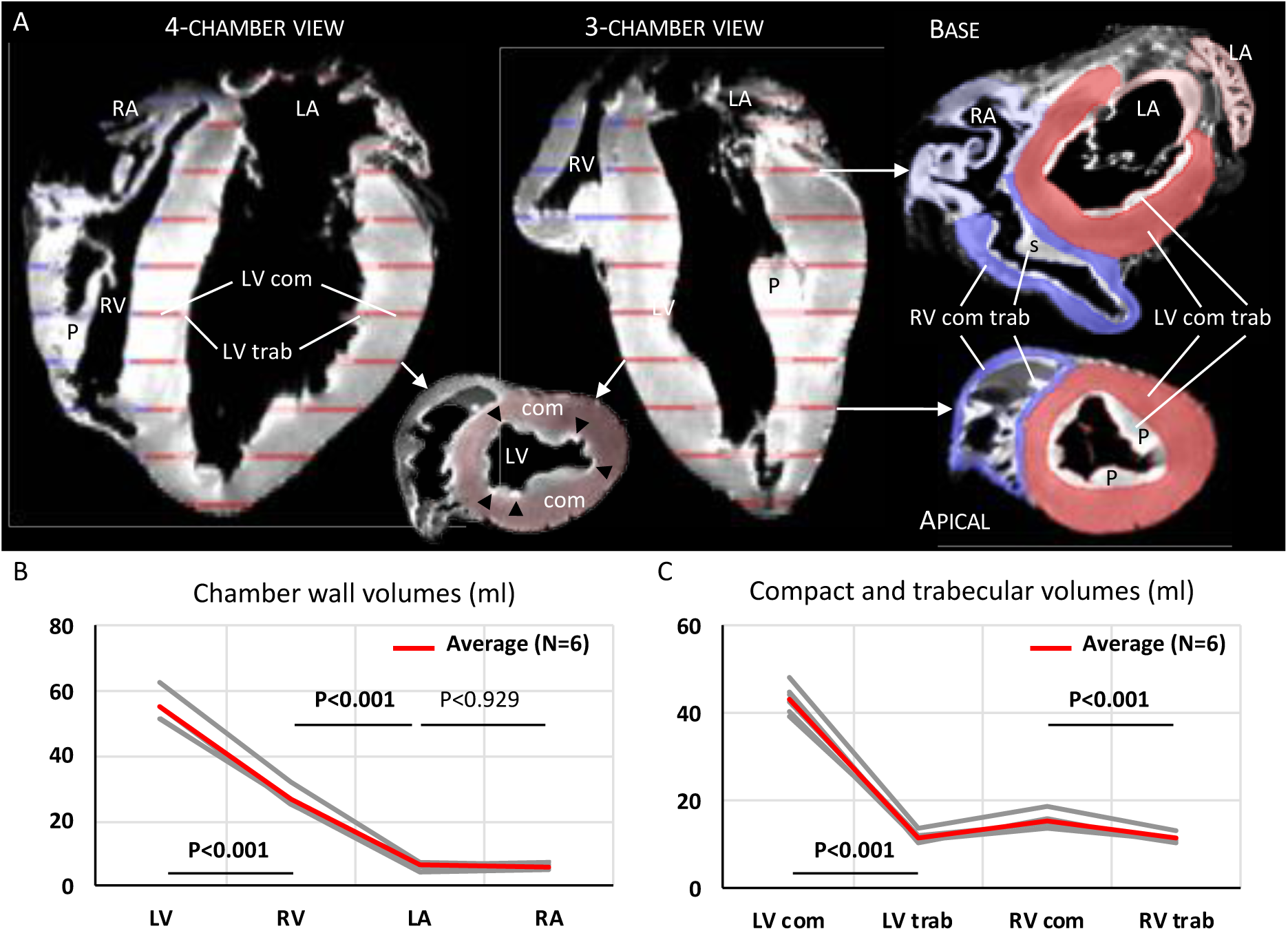
Volumetrics of the porcine heart. **A**. Every 10 transverse section was segmented for left ventricle (LV) and right ventricle (RV) trabecular (trab) and compact (com) wall, and left and right atrium. Notice most of the ventricular septum is segmented as being compact left ventricle. Structures that were prominent in the ventricular cavity were labelled as part of the trabecular layer, including papillary muscles (P) and most of the supraventricular crest of the right ventricle (s). In the small insert, black arrowheads points to blood coagulates in the deepest parts of the intertrabecular recesses, i.e. landmarks for identifying the boundary between compact wall and the trabecular layer. **B**. The left ventricle had the greatest tissue volume of the four chambers, the right ventricle an intermediate volume, and the two atria had the smallest volumes and these were not different from each other. **C**. Both the left and the right ventricle had significantly more compact than trabecular tissue. All tests were paired t-tests assuming unequal variance.

Myocardial volumes of the six human hearts was estimates in the same way as for the porcine hearts (Figure 6A-B). Total myocardial volume was 178.6 ml (±32.8, SD). As expected, the left ventricle had the most myocardium, followed by the right ventricle, and the volume of the left and right atrial walls were smaller than that of the right ventricle (Figure 6B). The fraction of atrial to ventricular myocardium was 0.22 (±0.06, SD), while the fraction of right atrium to left atrium was 1.07 (±0.17, SD) and not different from 1 (one sample t-test for difference from 1, P=0.383). Only the left ventricle had significantly more compact than trabecular myocardium (Figure 6C) and the left ventricle was proportionally less trabeculated than was the right ventricle (22.8% ± 1.8 versus 49.3% ± 2.1, t-test, P<0.001). The proportion of trabeculation of the left ventricle was not correlated to that of the right ventricle (P=0.362).

**Figure 6.**
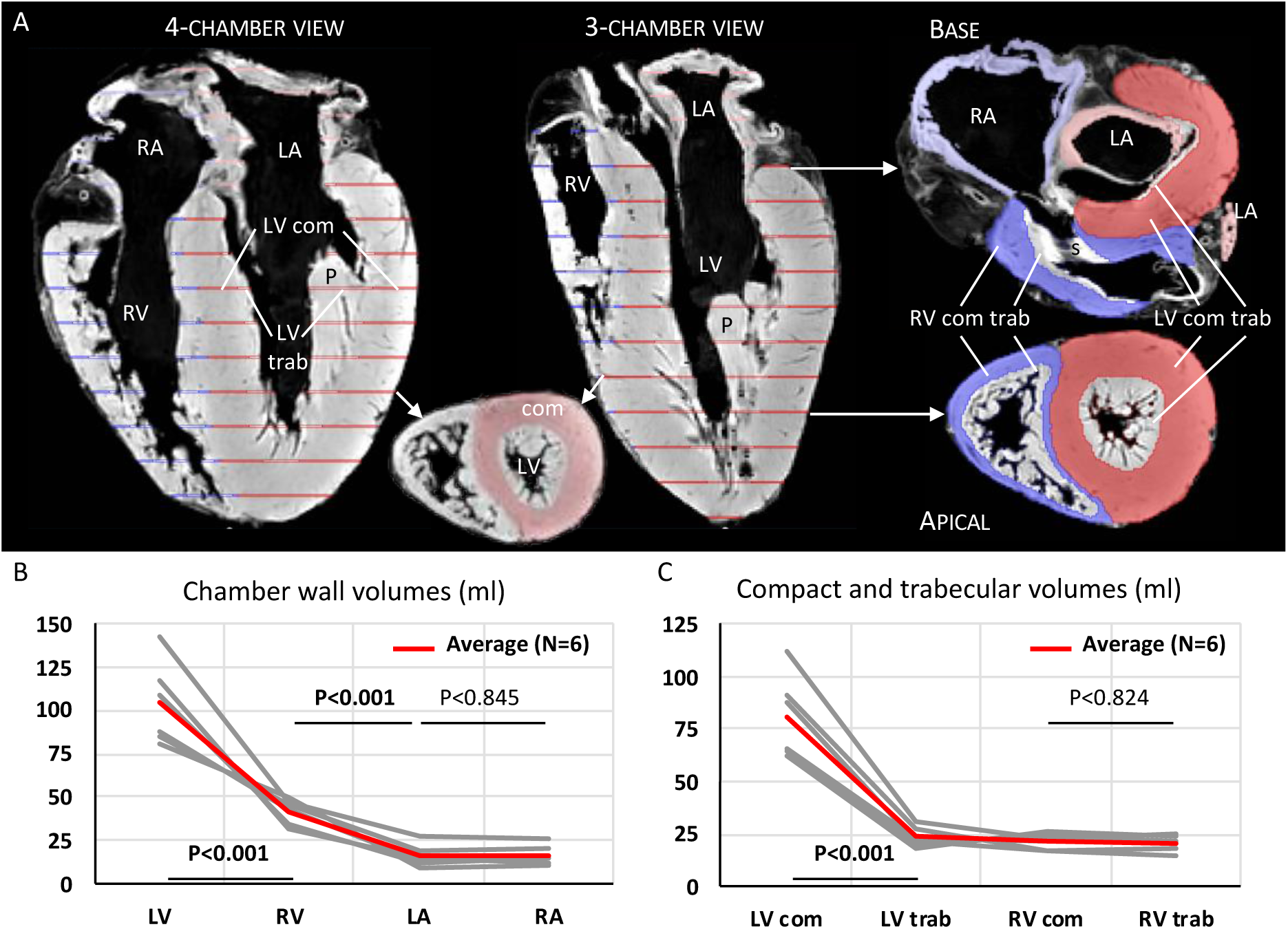
Volumetrics of the human heart. **A**. Every 15^th^transverse section was segmented for left ventricle (LV) and right ventricle (RV) trabecular (trab) and compact (com) wall, and left and right atrium. Structures that were prominent in the ventricular cavity were labelled as part of the trabecular layer, including papillary muscles (P) and most of the supraventricular crest of the right ventricle (s). **B**. The left ventricle had the greatest tissue volume of the four chambers, the right ventricle an intermediate volume, and the two atria had the smallest volumes and these were not different from each other. **C**. Only the left ventricle had significantly more compact than trabecular tissue. All tests were t-tests assuming unequal variance.

Given the almost twice greater myocardial volume of the human hearts relative to the porcine hearts, we only compared the proportions of trabeculation. The porcine right ventricle was less trabeculated than the human one (42.1% ± 1.4 versus 49.3% ± 2.1, t-test, P<0.001), whereas the porcine left ventricle was no less trabeculated than the human one (21.5% ± 1.7 versus 22.8% ± 1.8, t-test, P=0.240).

### Morphometrics of pig heart development

Myocardial and cavity volumes were measured in embryos from 14 days of gestation, at which point the chambers are only just forming, to 35 days of gestation, which, when compared with murine and human development, is well into the fetal stages of heart development (Figure 7A-C). The heart grows fast, at every older stage the heart is bigger, and between 14 and 35 days the ventricles increase approximately 400-fold in myocardial volume, from less than 0.001 ml to almost 0.4 ml (Figure 7D). Only the myocardial outflow tract, a tubular component downstream of the ventricles, decreased in length and tissue volume in the oldest stages (Figure 7D). Ventricular trabeculation was greater in volume in all older stages, in contrast to what could be expected if compaction transfers trabeculations to the compact wall (Figure 7D). In fact, for both ventricles the trabecular and compact tissue volumes were positively and significantly correlated (RV R^2^=0.97, P<0.001; LV R^2^=0.93, P<0.001; Figure 7E-F). Likewise, the thickness of the left ventricular compact wall increased approximately ten-fold (into more than 250 µm). Finally, while compaction supposedly would reduce the intertrabecular recesses, the recesses of both ventricles increased in volume to more than 0.01 ml, corresponding to an approximately 200-fold increase (Figure 7G).

**Figure 7.**
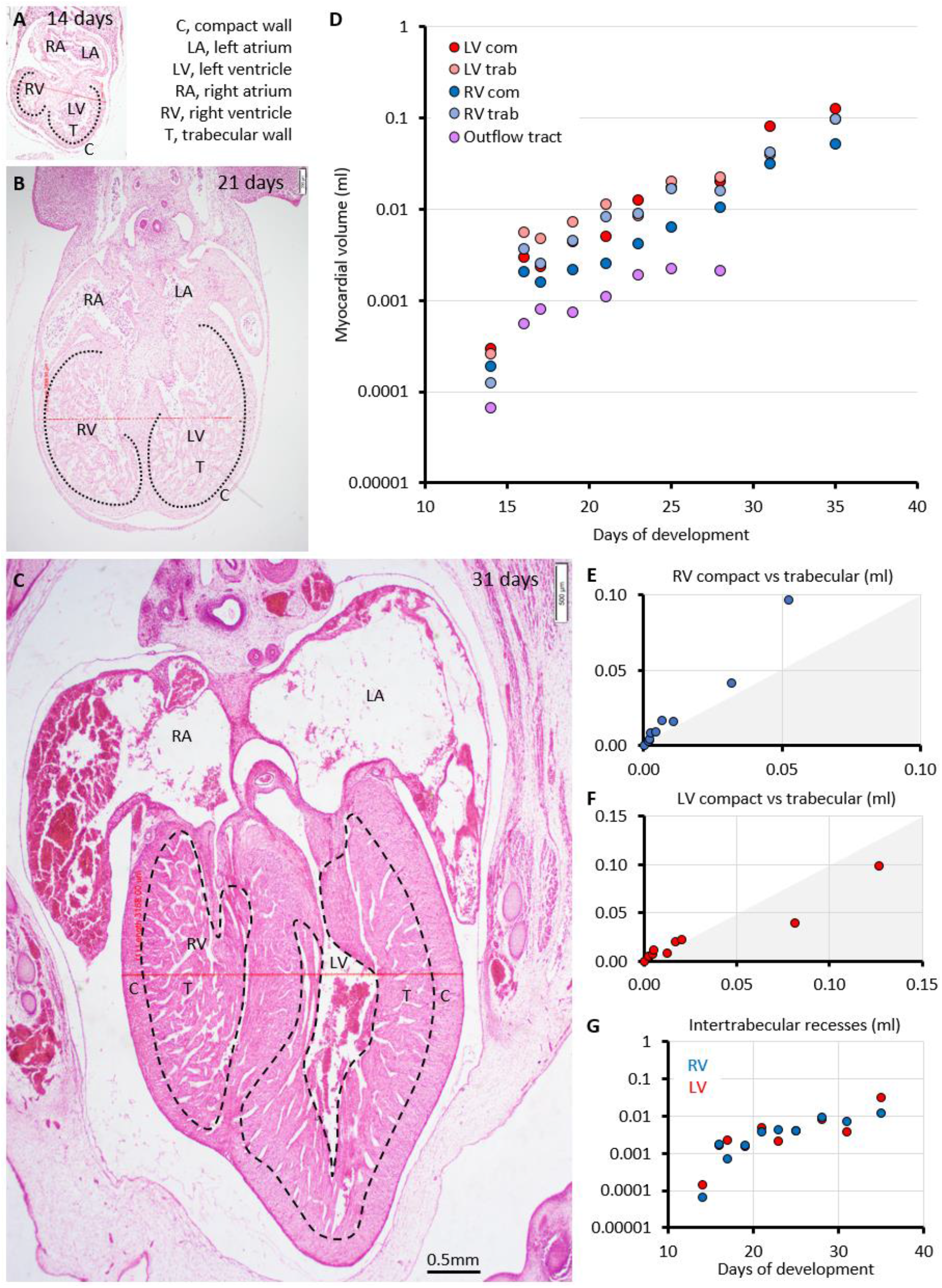
Pig heart development. **A**-**C**. Four-chamber view of developing hearts, on the same scale from gestational days 14, 21, and 31. **D**. All chamber components grow fast (the myocardial volume is displayed on a logarithmic scale), whereas the outflow tract did not, as expected [12, 51]. **E**-**F**. The trabecular and compact tissue volume was positively and significantly correlated for the right ventricle (**E**; R^2^=0.97, P<0.001) and left ventricle (**F**; R^2^=0.93, P<0.001). The shaded area indicates where more than 50% of myocardium is compact. **G**. The intertrabecular recesses of both ventricles increased much in volume.

Subsequently, we expressed the thickness of the trabecular layer as a ratio relative to the thickness of the compact layer, which is a ubiquitously used diagnostic criterion [42]. There was a rapid change from a ratio of approximately 3 at two weeks of gestation, to approximately 8 at around three weeks, and then down to 3 again in the oldest stages (Figure 8). The pronounced and fast change in the diagnostic criterion is then the outcome of the two layers growing at different rates rather than trabeculations becoming compact wall (Figure 8). The trabecular proportion of both ventricles peaked at around three weeks of gestation at which time approximately two-thirds of the ventricular myocardium was trabecular in organization. Interestingly, the adult proportions were not achieved even in the oldest stages in which much of the ventricular wall still had a trabecular-like appearance (Figure 9). Altogether, diagnostic measurements of non-compaction exhibit substantial changes in early heart development but they do not coincide with a decrement in trabecular myocardium.

**Figure 8.**
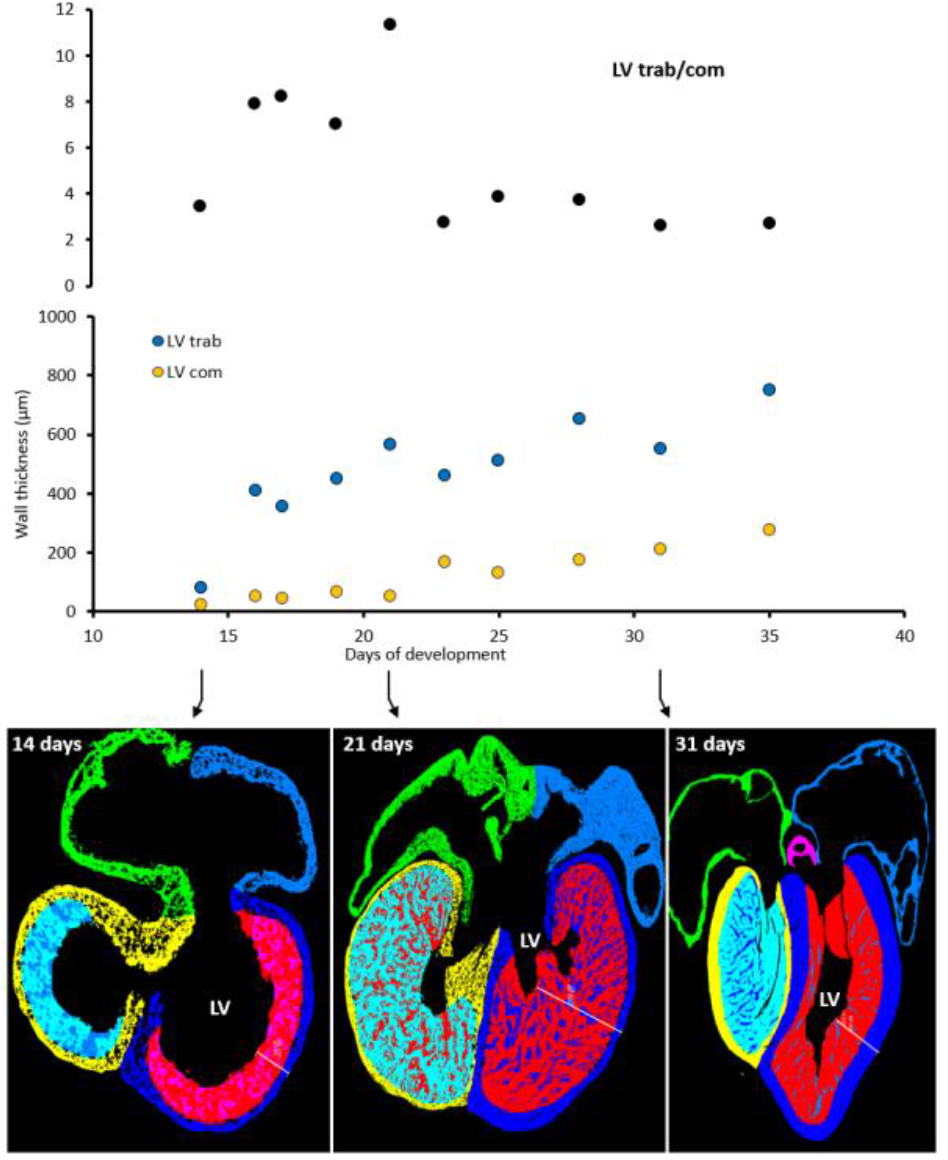
Differential growth rates drive changes to the relative thickness of trabecular layer. The left ventricular trabecular layer thickness relative to the neighboring compact wall thickness (LV trab/com) changes dramatically around 15-20 days and this change is driven by a faster thickening of the compact wall relative to the increasing widening of the trabecular layer. Architecturally, the 31 days LV wall looks much different from the proportionally much trabeculated LV of 21 days. Yet the trabecular layer of those two ages is approximately equally thick (measurements were done along the transmural line), whereas the older stage has an expanded cavity and a thicker compact wall which then gives the impression of a flattened of the trabecular layer which in fact it is not.

**Figure 9.**
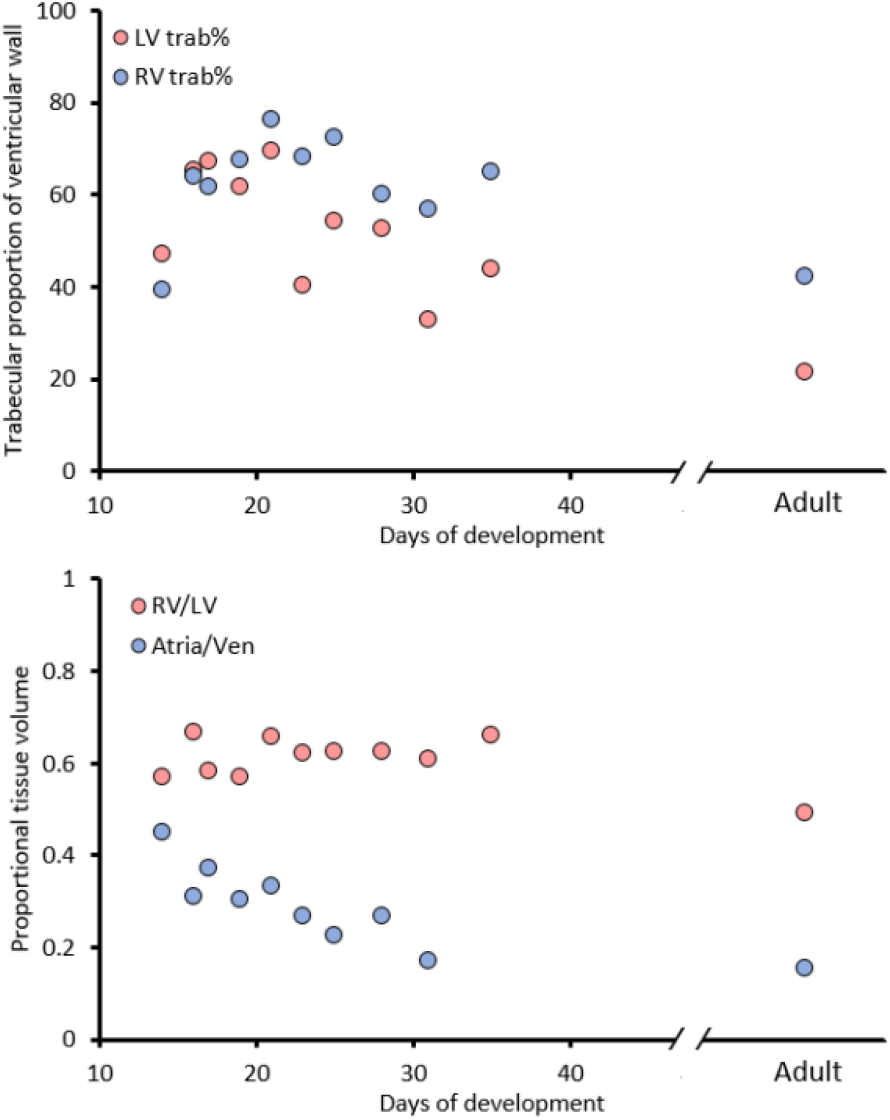
Proportional changes to chambers in pig heart development. The upper graph shows that the left and right ventricle (LV and RV respectively) are proportionally most trabeculated around 20 days. The adult proportions of trabecular muscle where not achieved even in the oldest stages. The lower graph shows that the LV is consistently bigger than the RV in early gestation. The atria progressively become smaller relative to the ventricles as in other vertebrates [51].

## Discussion

Visually, the pig ventricular walls could be the proof-positive result of extensive compaction because they clearly harbor many fewer trabeculations than do the equivalent walls of the human heart. Interestingly, the pig heart also has fewer trabeculations behind, for example, the papillary muscles of the right ventricle and such regions are located within the trabecular layer rather than being juxtaposed to the compact wall. The lower number of trabeculations there cannot be caused by trabeculations becoming compact wall i.e. by compaction. Perhaps the pig embryonic ventricle was never much trabeculated, but as shown before [52] and confirmed here, the porcine embryonic ventricles are much trabeculated. Rather than invoking compaction, the lower number of trabeculations in the adult heart can easily be explained if trabeculations coalesce with one another [13]. Coalescence could also easily explain trabeculations such as those numerated in Figures 1 and 4, which are several centimeters long and whose shape cannot be traced back to when the trabeculations first form in the embryo. Together, these observations suggest that a low number of trabeculations does not necessarily reflect the transfer of trabeculations to the compact wall, i.e. compaction. A process of trabecular coalescence within the trabecular layer appears to be a more likely explanation for the observations we report here.

An important challenge when assessing compaction from overview macroscopic inspection is revealed by close inspection and histology, namely that the porcine ventricular walls contain quite a few tiny trabeculations (many of these could very well be part of the Purkinje network). While these trabeculations are very small in the context of the adult heart, they are in fact bigger than the trabeculations of the embryonic heart that they have developed from. It is easy to miss the tiny trabeculations and if they are missed, the actual number of trabeculations will be underestimated. To the extent that a lower number of trabeculations equates greater compaction, then, the effect of compaction in pig is easy to overestimate.

Early heart development in pigs and humans are quite similar in the sequence of appearance of structures [53]. Two fairly detailed accounts of the development of the porcine ventricular walls are published [10, 52], but they appear to be contradictory. In the paper by Shaner [52], a zone of trabeculation-like “ridges” is described to develop from the compact wall, or “cortical” layer. These ridges are located between the trabecular, or “sponge”, layer and the compact wall and they have seemingly formed by a sort of de- compaction. Our analysis did not confirm or disprove the presence of such process. The other study, by Gabriel and colleagues [10], emphasizes a developmental thickening of the compact wall brought upon by compaction around days 26 and 30 even though actual thicknesses are not reported and the scale of the supporting images are not given. In our series, the greatest decrement in the ratio of trabecular-compact wall thickness occurs in the days immediately before day 26. In our view, this decrement may equate to what Gabriel and colleagues [10] call compaction. But the decrement is brought about by a greater thickening of the compact layer compared to the thickening of the trabecular layer, in absolute terms, rather than by compaction.

Here, we quantified the trabecular and compact layers by actual volumes, proportions, and thicknesses in addition to the volume of intertrabecular recesses. While compaction could be expected to entail a decrease of the trabecular layer thickness and volume, in addition to disappearance of the intertrabecular recesses, all of these measurements instead showed an increase. In this way, pig heart development resembles early heart development of humans, mice, shrews, and chicken [12, 21, 54]. Morphometry, then, does not support a substantial role for compaction in determining the morphology of the porcine ventricular wall. The formed pig heart, nonetheless, macroscopically appears much more compacted than does the adult human heart. This state of greater compaction, however, appears to be a misinterpretation, because the proportion of trabecular myocardium in the formed heart is not much different in pigs and humans. In essence, the porcine ventricles compared with the human ventricles are dominated macroscopically by fewer but larger trabeculations, while the proportional volumes of the trabecular and compact layers are quite similar between the two species. In pig, then, it is likely that coalescence of trabeculations changes the trabecular layer the most. The effect of coalescence on the compact layer is more subtle if present at all. To what extent coalescence is involved in the formation of large atrial trabeculations, the pectinate muscles, is not clear at this point.

Perhaps the strongest case for substantial compaction is the observation of Purkinje cells deep within the formed compact wall which is found in pigs and in ungulates in general [55–58]. Studies in mouse, including lineage tracing, show that the peripheral Purkinje system originates from the embryonic trabecular layer [14, 16, 59]. In mouse, and human, the Purkinje system is only on the endocardial side of the wall and it does not have the deep intramural penetrance seen in ungulates. The simplest assumption must be that in pig the Purkinje cells originate from the embryonic layer only, which in turn suggests substantial compaction because of the deep penetrance of the Purkinje cells, but that assumption has not been tested. Although our developmental findings cover the entire period in which compaction is said to occur [10], the ventricular wall of the oldest periods still appears immature compared to the ventricular wall morphology of the adult animal. Specifically, there are still relatively deep intertrabecular recesses and the trabecular proportion has not yet reached the low levels of the adult heart.

The transition from a proportionally much trabeculated ventricle to one dominated by the compact wall is seen in normal development of the endothermic mammals and birds. This development is different from that of hearts of reptiles which likely represents the evolutionary ancestral condition to hearts of mammals and birds [4, 60–63]. In reptiles, the highly trabeculated state is maintained. Not much has been done to understand the evolutionary transition from the highly trabeculated wall to one dominated by compact myocardium. It is speculated to relate to a reduction in the time it takes to fill and empty the ventricles and this could enable high heart rates [36, 64, 65].

The developmental transition from the highly trabeculated wall to one dominated by compact myocardium has mostly been investigated in a biomedical context to give perspectives to so-called noncompaction [37, 61, 66–68]. Since noncompaction presupposes compaction, the increasing interest in noncompaction [17, 27, 69] has likely led to a misattributed and exaggerated importance to compaction [13, 18, 46]. In biology, shape change is often driven by differential growth rates [70], and this is also the case for the pig, as we show here, and in human, mice, and chicken [21], and even in animals with an almost ‘smooth’ ventricular wall such as shrews [54].

### Limitations

Compared to the humans whose hearts were studied, the pigs had experienced a lot less variation in age and they were comparatively young. At least in human, the degree of trabeculation may be under some influence of life history. Compared to humans, pigs raised for slaughter grow much faster. It is not clear to us, to what extent these factors exaggerate or diminish the well-established species differences in trabeculation [9].

Concerning the precision of the morphometric measurements based on MRI and to a lesser degree also on histology, the discrimination of ventricular trabeculations from lumen appears good, especially when blood coagulates are absent. In contrast, the discrimination from lumen of even the largest atrial trabeculations, the pectinate muscles, is difficult and between these there can be numerous finer trabeculations. In addition, it is challenging to define the border between trabeculations and compact wall in a precise and standardized manner. These imaging limitations, together the expected 5-10% error in volume estimates per specimen, may ultimately have obscured small differences in trabeculation between pig and human.

## Conclusion

Our volumetric analysis of porcine heart development shows that compact and trabecular myocardium grows and cavities expand. These findings do not support a role for compaction in shaping the ventricular wall architecture. The observations on adult hearts of strikingly fewer trabeculations in pig compared to human could suggest extensive compaction in pig. It is then somewhat contradictory, however, that the proportions of ventricular trabecular myocardium are much similar in pig and human. There is no contradiction if, in pig, many and small trabeculations are coalesced to fewer and larger trabeculations. Such process of coalescence is likely restricted to the papillary muscles in human, whereas in pig it could affect the trabecular layer at large.

## Acknowledgements

The authors have no conflicts of interest to report.

